# How predator hunting-modes affect prey behaviour: Capture deterrence in *Drosophila melanogaster*

**DOI:** 10.1101/010330

**Authors:** Abhijna Parigi, Cody Porter, Megan Cermak, William R. Pitchers, Ian Dworkin

**Author notes:** Correspondence to Ian Dworkin.

## Abstract

Hunting mode, the distinct set of behavioural strategies that a predator employs while hunting, can be an important determinant of the prey organism’s behavioural response. However, few studies have considered how a predator’s hunting mode influences anti-predatory behaviours of a prey species. Here we document the influence of active hunters (zebra jumping spiders, *Salticus scenicus*) and ambush predators (Chinese praying mantids, *Tenodera aridifolia sinensis*) on the capture deterrence anti-predatory behavioural repertoire of the model organism, *Drosophila melanogaster*. We hypothesized that *D. melanogaster* would reduce overall locomotory activity in the presence of ambush predators, and increase activity with active hunters. First we observed and described the behavioural repertoire of *D. melanogaster* in the presence of the predators. We documented three previously undescribed behaviours-abdominal lifting, stopping and retreat-which were performed at higher frequency by *D. melanogaster* in the presence of predators, and may aid in capture deterrence. Consistent with our predictions, we observed an increase in the overall activity of *D. melanogaster* in the presence of jumping spiders (active hunter). However, counter to our prediction, mantids (ambush hunter) had only a modest influence on activity. We also observed considerable intra and inter-individual variation in response to both predator types. Given these new insights into *Drosophila* behaviour, and with the genetic tools available, dissecting the molecular mechanisms of anti-predator behaviours may now be feasible in this system.

## INTRODUCTION

Predation, a ubiquitous selective force, gives rise to and determines the nature of defensive traits in prey populations (Edmunds, 1974; Juliano & Gravel 2002; Goslin & Rodd, 2007; Langerhans, 2007; Lima & Dill, 1990; Sansom, Lind & Cresswell, 2009). Predator hunting-modes, i.e., the set of behaviours that predators employ to pursue and capture their prey (Schoener, 1971; Huey & Pianka 1981; Preisser, Orrock & Schmitz 2007), have been shown to induce distinct prey responses (Schmitz, 2008) that in turn influence the productivity of ecological communities. In habitats dominated by active hunters there is lower species evenness and higher above-ground net primary productivity compared to habitats dominated by ambush hunters (Schmitz, 2008). The authors suggest the observed differences in prey productivity to be driven by hunting mode specific trade-offs between foraging and seeking refuge. Although studies often describe the effects of predators on prey traits (i.e. DeWitt, Robinson & Wilson, 2000; Reznick, Butler & Rodd, 2001; Relyea, 2001), it is rare for the role of predator hunting-mode to be explicitly considered.

Here we investigate segregating differences in the anti-predatory behavioural repertoire of the fruit fly, *Drosophila melanogaster*, in response to two predator species differing in hunting modes. Based on (Schmitz, 2008), we predicted that fruit flies, in the presence of a familiar predator, would exhibit hunting-mode specific modifications in activity levels. We used *D. melanogaster* because, although it is one of the most well-studied model organisms, there is a relative paucity of information regarding *D. melanogaster’s* natural history, ecology and behaviour, including habitat, food resources, and natural enemies (but see Reaume & Sokolowski, 2006; David & Capy, 1988; Turelli & Hoffmann, 1991; Schmidt, et al. 2005; Fleury et al. 2004; Wilfert & Jiggins, 2014; Stephan & Li, 2006). While anti-predator behaviours are well studied as targets of selection in prey (Juliano & Gravel, 2002; Stoks, McPeek & Mitchell, 2003; Magurran et al. 1992), the genetic bases of such behaviours are seldom investigated. Given the range of genetic and genomic tools available for *D. melanogaster,* along with its complex behavioural repertoire and suitability for experimental evolution, understanding the anti-predatory behaviours persisting in a natural population of the fruit fly brings us one step closer to deciphering the molecular mechanisms for anti-predator behaviours.

Previous work has examined the effects of natural enemies on population and community structures of *Drosophila spp*. Worthen (1989) studied the effects of predation by rove beetles (staphylinids) on the coexistence of three mushroom-feeding *Drosophila* species, and Escalante & Benado (1990) showed that ant predators regulate population densities of wild *D. starmeri* (cactophillic fruit fly). In *D. melanogaster per se*, the role of parasites in influencing larval and adult behaviours has been extensively studied (Milan, Kacsoh & Schlenke, 2012; Kacsoh et al., 2013; Polak & Starmer, 1998). Despite this literature, we know little about the predators of *D. melanogaster* adults in the wild, nor the nature of anti-predatory behaviours segregating in natural populations.

We documented the influence of two predators, the zebra jumping spider (*Salticus scenicus*) and juvenile Chinese praying mantids (*Tenodera aridifolia sinensis*) on the capture-deterrence behaviours of *D. melanogaster* individuals derived from a wild-caught population. The zebra spider is an active hunter, locating prey visually (with an extensive visual field attained by antero-medially positioned simple eyes) (Dill, 1975; Horner, Stangl & Fuller, 1988). Mantids are generally ambush predators, waiting for prey to enter their attack range (Prete, Klimek & Grossman, 1990). Despite numerous differences, zebra spiders and juvenile Chinese mantids are similar in two relevant ways. First, both species primarily detect prey visually (Forster, 1979; Harland, Jackson, Macnab, 1999; Jackson, & Blest, 1982; Prete 1999) and are likely incapable of depth perception when their prey item is motionless (Prete, 1999; Freed, 1984). Second, small adult diptera account form a substantial proportion of the diet of both predators in the wild (Iwasaki, 1998; Okuyama, 2007).

Based on the findings of Schmitz (2008), we predicted that fruit flies, in the presence of a familiar predator, would exhibit hunting-mode specific modifications in activity levels. To maximize distance from the actively hunting spider, our prediction was that flies would increase their overall activity levels, whereas, to reduce the probability of encountering a stationary threat (the mantid, an ambush predator), we expected flies to decrease overall activity.

Under controlled laboratory conditions, we documented the behaviours of individual adult *D. melanogaster* with and without the two predator species. Our results suggest that in the presence of zebra spiders, *D. melanogaster* increases its overall locomotory activity, performs a distinct “stopping” behaviour and increases the performance of a newly described abdominal lifting behaviour (the function of which is as of yet unknown). Counter to our prediction though, *D. melanogaste*r’s locomotion, and most other behaviours are not substantially altered in the presence of mantids. However, upon direct encounter with a mantid, many individuals of *D. melanogaster* perform (a previously undescribed) retreat behaviour- a response not generally elicited by jumping spiders. Furthermore we observe considerable intra- and inter-individual variation in response to predators. We discuss our results in terms of conditionally expressed behaviours as they relate to predator hunting mode, co-evolutionary history of predators and prey, and in terms of broadening our understanding of the behavioural ecology of *D. melanogaster*.

## METHODS

### Drosophila Population and Culture Conditions

The *Drosophila melanogaster* population used in this study originated from a natural population at Fenn Valley Vineyards in Fennville, Michigan (GPS coordinates: 42.57, -86.14) during the summer of 2010. A lab population (henceforth referred to as FVW) was initiated from this collection using the progeny of over 500 single-pair matings of field caught *D. melanogaster* as well as wild caught males. This design allowed us to screen out the sympatric congener, *D. simulans,* which was present in our collections at a frequency of about 5%. Screening involved setting up single pair mating in vials and discarding all lines with *D. simulans*-like genital morphology. After screening, ∼1500 individuals were placed into cage (32.5cm^3^, BugDorm BD43030F) to establish the FVW population. The population is currently maintained in this cage at an adult density ∼ 3000 individuals in a room maintained at 23°C (+/-1°C), and 40-70% RH. Adults were allowed to lay eggs in 10 bottles with 50-60 ml of a standard yeast-cornmeal food for 2-3 days. These bottles were then removed and kept in a Percival incubator (Model: I41VLC8) at 24°C and 65% RH throughout the larval stages. All flies and larvae were maintained in a 12 hr light/dark cycle with lights on at 08:00 hours.

For the experiments, pupae were collected 24 hours before they emerged as adults. Pupae were removed from bottles using forceps and individual pupae were placed into 1.5 ml microcentrifuge tubes. Each tube was pre-filled with ∼ 0.5 ml of yeast-cornmeal food and its cap was punctured for gas exchange. Upon emergence, adult flies were sexed visually without anesthesia and housed in these tubes in the incubator until needed for behavioural assays. Age of flies used in behaviour analysis was 3-7 days. By using socially naïve flies in our assays, we were able to establish a consistent baseline of social experience among all individuals, allowing us to eliminate the potentially confounding influence of variation in social experience on behaviour that is well-documented in *Drosophila* (Yurkovic et al., 2006; Levine, 2004; Krupp et al., 2008; Lefranc et al., 2001; Chabaud et al., 2009).

### Spiders

*S. scenicus* individuals were collected throughout the spring/summer of 2012 on the campus of Michigan State University. Spiders were housed individually in vials in a room maintained at 23°C (+/-1°C) and 30-50% RH and fed ∼5 *D. melanogaster* a week. Prior to use in behavioural assays, spiders were starved for at least 48 hours. Each spider was used in only a single behavioural assay.

### Mantids

Mantid egg cases were both collected near the campus of Michigan State University as well as ordered from Nature’s Control (Medford, Oregon). Mantid egg cases were stored at 4°C and transferred to 25°C and 70% RH for hatching. Given the substantial changes in mantid body size across moults (Iwasaki, 1990), only first instar nymphs were used for experiments. Prior to behavioural assay, mantids were starved for at least 24 hours and each mantid was used only once.

### Behavioural Assays

All assays were performed 1-4 hours after the incubator lights came on in the morning (08:00). Behavioural assays were recorded with an Aiptek AHD H23 digital camcorder attached to a tripod under a combination of natural and fluorescent light that is present in the room wherein the FVW population and spiders are maintained. For each predator (spiders and mantids), we recorded the behaviour for each of 15 male and 15 female socially naïve, virgin flies (collected as described above). We used a chamber constructed from the bottom of a 100 ×15mm petri dish inverted on top of a glass plate with a sheet of white paper beneath to maximize the visibility of flies and predators.

For each assay, an individual fly was aspirated into the chamber and allowed to acclimate for 5 minutes. After this acclimation period, flies were recorded for 5 minutes. A single spider or mantid was then introduced to the chamber and behaviours were recorded for an additional 10 minutes or until capture. The chamber was washed with 10-30% ethanol and rinsed with reverse osmosis water after each assay to remove olfactory cues.

### Behaviours Recorded

All *Drosophila* behaviours were categorized and analysed as either “states” or “events”. Behavioural states have measurable duration and are mutually exclusive with other states (e.g. individuals cannot simultaneously walk and run). Behavioural events are discrete behaviours that occur instantaneously and are also mutually exclusive with each other (e.g. turning versus jumping) but not always mutually exclusive with behavioural states. For example, an individual could perform a wing display (event) while simultaneously walking (state), but it could not jump (event) while simultaneously running (state). In this study we treated flying as an event because the structure of the experimental chamber constrained flight duration. Attempted flight by *D. melanogaster* could result in landing due to contact with a wall of the petri dish. We also recorded when a fly was not visible (occluded) to the observers analysing video. We recorded a total of 6 discrete events and 5 behavioural states in *D. melanogaster* in response to predation by spiders and mantids (Table 1). In order to interpret an individual fly’s behaviour in the context of predatory encounters, we designated two keys to describe the location of the predator in regard to its interactions with the fly. As flies might alter their behaviour when a predator is within striking distance, we recorded predator location based on whether or not it was within striking distance of the fly (∼ 5mm from the spider/mantid, also see *Spider location*/ *Mantid location* in Figure 1).

**Table 1.**
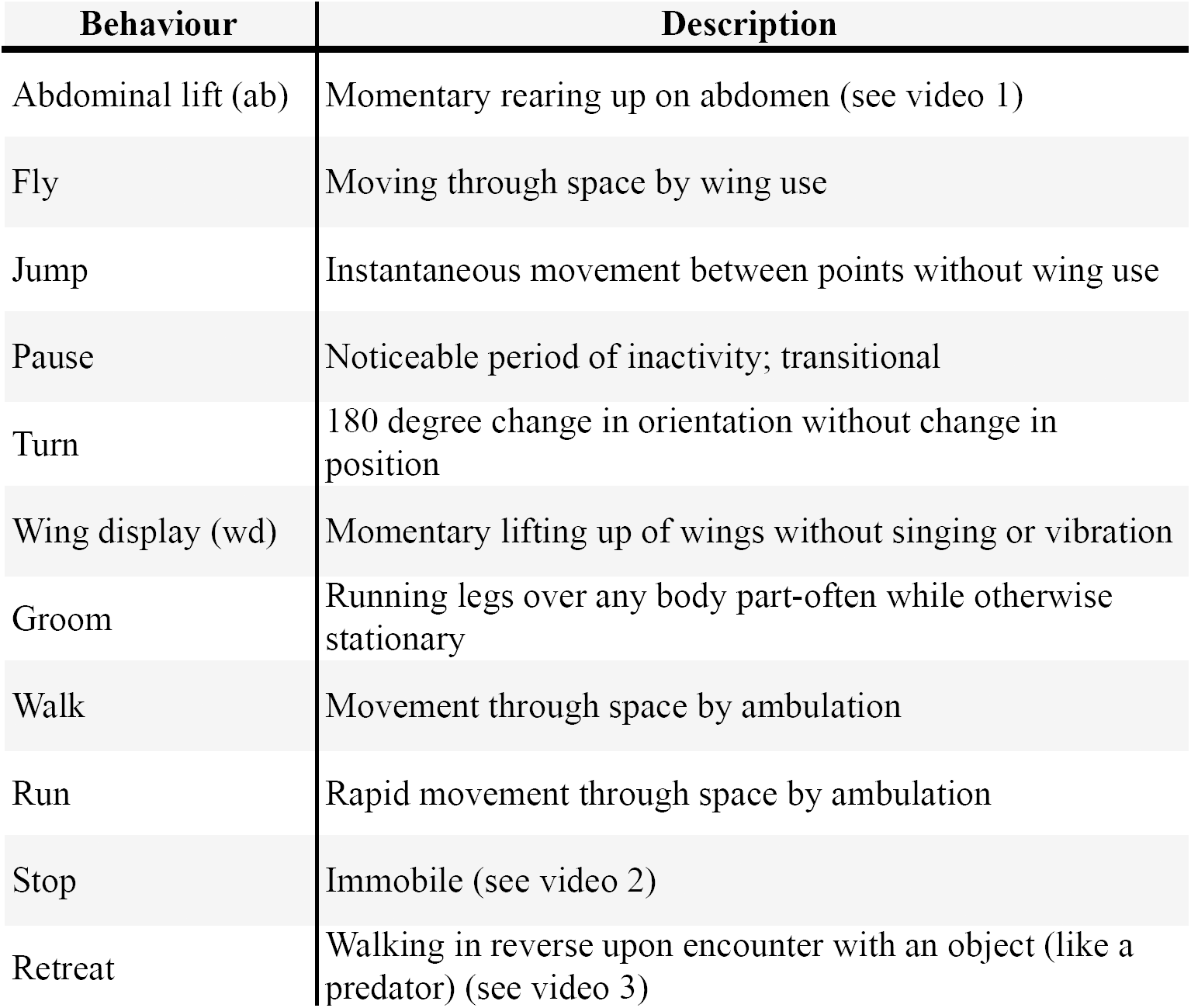
Names and descriptions of all observed behaviours. Videos are provided at the end of Supplement b.

**Figure 1.**
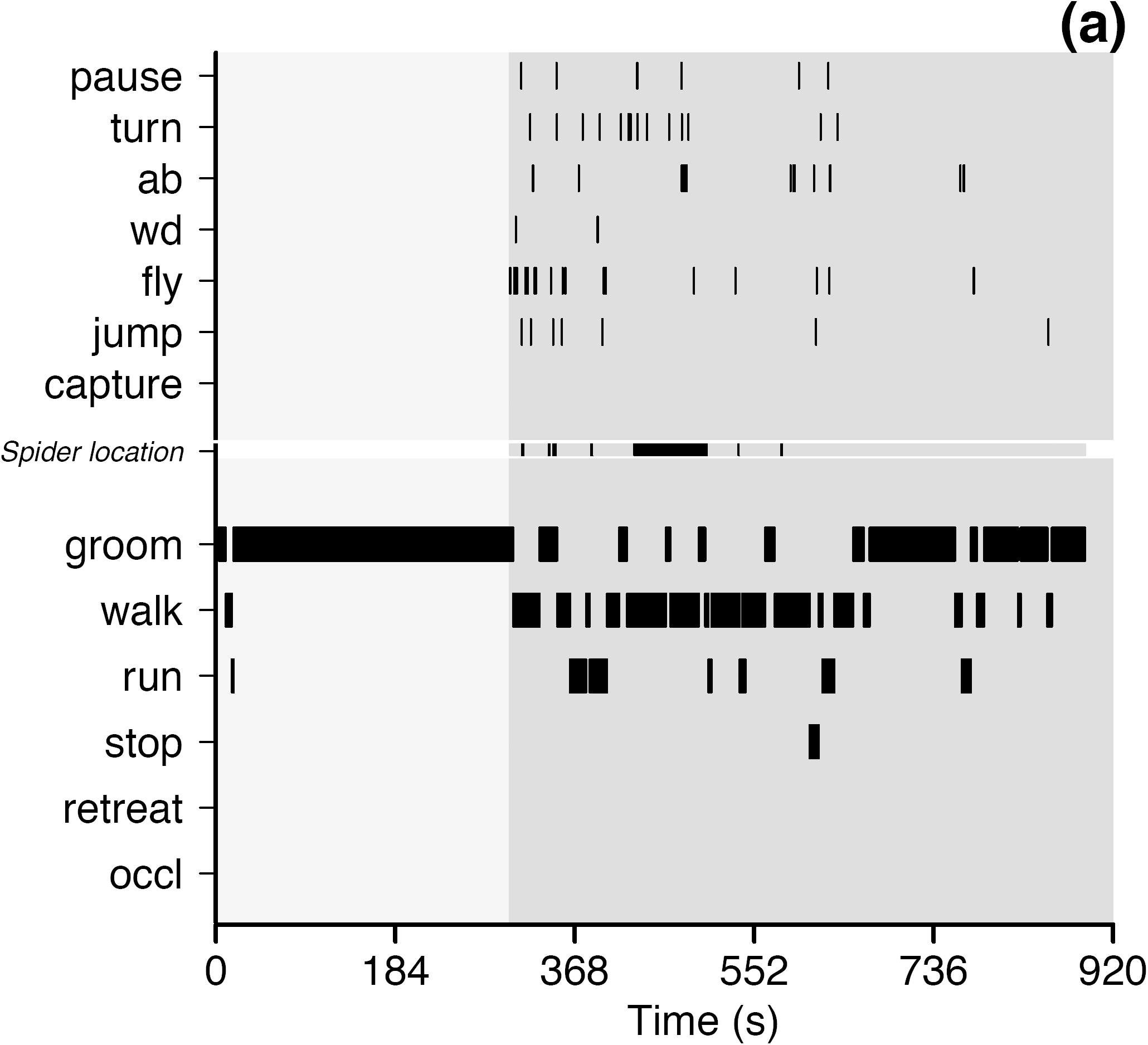
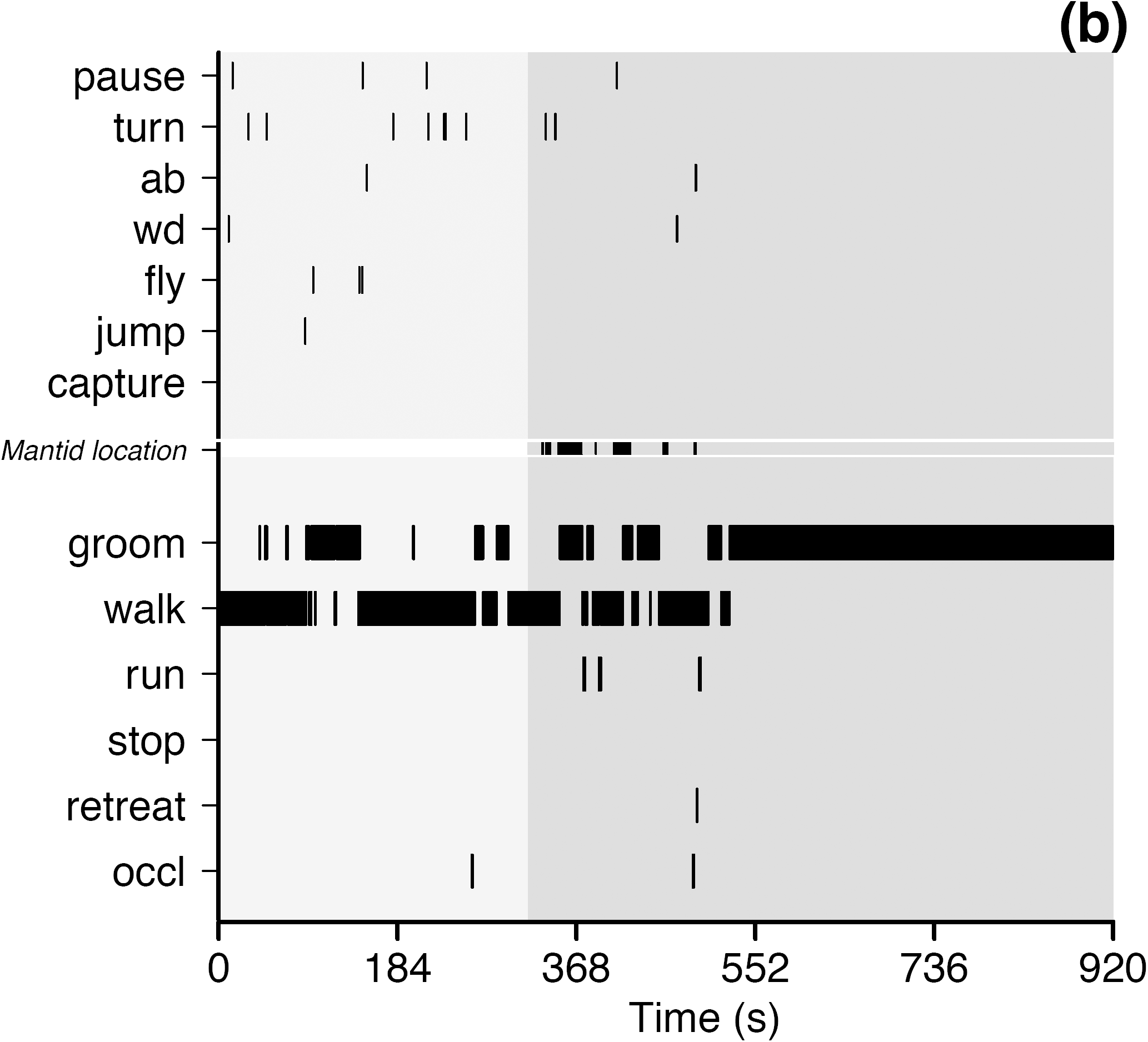
*D. melanogaster* used a greater proportion of its behavioural repertoire and performed each behaviour at a higher frequency in the presence of a jumping spider than in the presence of a juvenile mantid. a) Representative ethogram of a male, 4 day old *D. melanogaster* in response to a zebra jumping spider. b) Ethogram of a male, 5 day old *D. melanogaster* in response to a juvenile Chinese paying mantid. Light grey background represents time in the arena before the addition of a predator and dark grey background is when the predator was present in the chamber. Each black bar represents the occurrence of a behaviour during the observation period. Top half of the figure (separated by *Predator location*) consists of events and the bottom half consists of states. Because states have duration, the width of each black bar corresponds to the duration of a state. *Predator location* (i.e., *Spider location* in a and *Mantid location* in b) indicates whether or not the predator was within striking distance of the fruit fly at that time point. This information is relevant only after the predator was added to the chamber (∼ 300 s into the assay). Dark grey bars in *Predator location* indicate that the spider was within striking distance and light grey regions indicate that the spider was out of striking distance. *Predator location* is white when the predator is absent from the arena or after successful capture. If capture did not occur, *predator location* remains light grey in colour.

### Video Processing

Recorded behaviours were viewed with VLC media player (version 2.0.3) and analysed by two observers using a manual event recorder, JWatcher V1.0 software (Blumstein, 2006). One observer (A.P.) viewed each video and verbally announced the occurrence of behaviours while the other observer (C.P./ M.C.) recorded the occurrence of these behaviours with JWatcher. Because *Drosophila* anti-predatory behaviours are often complex and occur rapidly, we analysed all videos at 0.5X speed.

### Controlling for effects of season and disturbance

We conducted experiments with spiders between October and December 2012 and those with mantids from March and May 2013. To confirm that predator species-specific behavioural differences were not confounded with seasonal differences in behaviour, we performed 6 additional assays (alternating between spider and mantid treatments) within the span of one week. Following a spider assay, the plates were wiped down with 30% ethanol followed by a rinse with RO water before a mantid assay was conducted.

Additionally, the process of adding a predator to the arena invariably resulted in a disturbance that likely startled the fly (unrelated to the presence of a predator). To confirm that behaviours induced by this disturbance were not confounded with predator induced behavioural differences, we performed 3 control assays. Here, after 5 minutes of acclimatization without a predator (see above for more details), the arena containing the fruit fly was disturbed gently (∼ magnitude of disturbance caused by the addition of a predator). For all controls, video processing and behaviours recorded were identical to mantid and spider treatments described above. See Supplement b, S1 for a detailed description of these control experiments and their results.

### Data processing and statistical analysis

A custom Python script was used to parse Jwatcher formatted data files into a comma-separated-value (CSV) file for analysis in **R** (version 3.0.1).

To analyse the effects of predator state (i.e., presence or absence of predators) on the time dedicated to locomotory behavioural states, and number of occurrence for behavioural events, we fit generalized linear mixed effects models (using both `glmer` function in the lme4 package version 1.0-5, and the `MCMCglmm` function in the MCMCglmm package version 2.17) with predator state, total duration of assay with and without a predator (duration), sex, temperature and recording time as fixed effects, and individual by predator state and date as random effects. Formally, the model was:

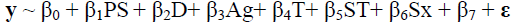

Where **y** is a vector of time spent in a behavioural state. β_1_ is the regression coefficient for predator state, β_2_ is for duration in each predator state, β_3_ is for age of the fly, β_4_ is for temperature, β_5_ is for time at which assay was started, β_6_ is for sex of the fly and β_7_ is for date on which the assay was performed. We estimated random effects for individuals including variation in response to predator state and duration of assay, and we fit an independent random effect for date. Thus we fit a repeated effects (longitudinal) mixed effects model allowing for variation among individuals for the influence of predator presence and duration of assay where for the i^th^ individual

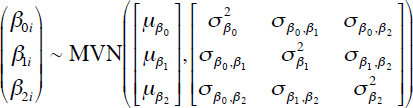

and (independent of the above)

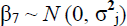 where j = 1 … date

Preliminary analyses were inconsistent with the need to fit higher order interactions among fixed effects, so interaction terms were not considered further. The one exception was for “stopping” behaviour where individuals almost exclusively performed this in the presence of the predators. For the behavioural states (locomotion, grooming and stopping) we assumed normally distributed variation. For the counts of events (abdominal lift, jumping, etc) we used a log-link function and assumed the variation was poisson distributed. Estimation using both maximum likelihood (`lmer`) and simulating the posterior distribution (`MCMCglmm`) provided similar results for fixed effects, and generally for random effect components as well.

Among individual coefficients of variation were calculated by dividing the square root of the among individual variance component from the model by its respective fixed effect estimate (i.e. its “mean”). While confidence intervals were consistent for fixed effects, the intervals were more difficult to estimate given the complexities of the random effect structure of the model, and some caution is warranted for their interpretation.

To test for non-random associations in the temporal structure of behavioural patterns we constructed transition frequencies using the “msm” library (version 1.2) (Jackson, 2011) in **R**. To test for both for first order Markov processes between behaviours (transition probabilities), as well as the influence of predator presence on these transition probabilities, we fit log-linear models (assuming poisson distributed data) with the transition frequency matrices (Crawley, 2012) using `glm` in **R**. As advocated by (Crawley, 2012; Bakeman & Gottman, 1997) we fit a saturated log-linear model (with lag0, lag1 and predator state as the effects in the model) and tested the influence of deleting the terms (i.e. third order interaction) on change in deviance. We used modified “Z-scores”, adjusted using sequential Bonferroni to assess the deviation of particular cells in the transition frequency matrix from expected values (assuming independence). For the visual transition probability matrices, we combined the behavioural event “pause” with the behavioural state “stop” because 1) we wanted to reduce the complexity of the matrix and 2) the main difference between the two behaviours is that pause is instantaneous and stop has duration. All transition diagrams were constructed in Inkscape (version 0.48.2, Harrington, 2004-2005).

## RESULTS

From pilot observations (not included in analysis), we (I.D., A.P. and C.P.) catalogued and described *Drosophila melanogaster* behaviours observed in the presence of a predator (Table 1). Among the behaviours listed in Table 1, <u>abdominal lifting</u> (ab, supplement b, video 1) and <u>retreat</u> (supplement b, video 3), to our knowledge, have not been previously described in *D. melanogaster* literature.

### Flies perform a range of anti-predatory behaviours in response to a zebra spider

To visualize each individual fruit fly’s response to the presence of a zebra jumping spider, we generated ethograms (see Figure 1a and Supplement a). For the two predator states (spider present and spider absent) we measured the mean proportion of time dedicated to each behavioural state, as well as the number of occurrences per minute for each behavioural event. When a spider was present, *D. melanogaster* increased the proportion of time it spent walking and running by 50% (95% CI: 21-79% increase) while grooming 60% less (95% CI: 43-77% decrease). This is shown in Figure 2 and Supplement b Figure S1 (treatment contrasts with 95% CI in figures are provided to enable assessment of significance). While they were observed at low frequencies prior to the addition of spiders, *D. melanogaster* substantially increased the frequency of pauses, jumps and flights (per minute) in the presence of spiders (Figure 2, Supplement b Figure S3). For instance the frequency of abdominal lifts increased from 1.51/minute to 4.0/minute (95% CI: 1.55-9.93), while jumping showed a 6.6X increase from 0.73/minute to 4.82/minute (95% CI: 1.77-12.17). “Stopping”; a motionless state that likely aids in capture deterrence (see Videos 2, Supplement b) was not performed by *D. melanogaster* in the absence of spiders (Supplement b Figure S2). However in the presence of spiders, the average total time spent “stopping” increased to ∼25.8 seconds (95% CI: 10.1 - 41.7 seconds). When interacting with spiders, flies were only observed to perform the “retreat” behaviour once (of 30 individuals). Interestingly, we did not see significant sex specific differences in either frequencies of occurrence (Supplement b Figure S3) or proportion of time allocate (Supplement b Figure S1) to the majority of measured behaviours (But see S3 panels “pause” and “turn”).

**Figure 2.**
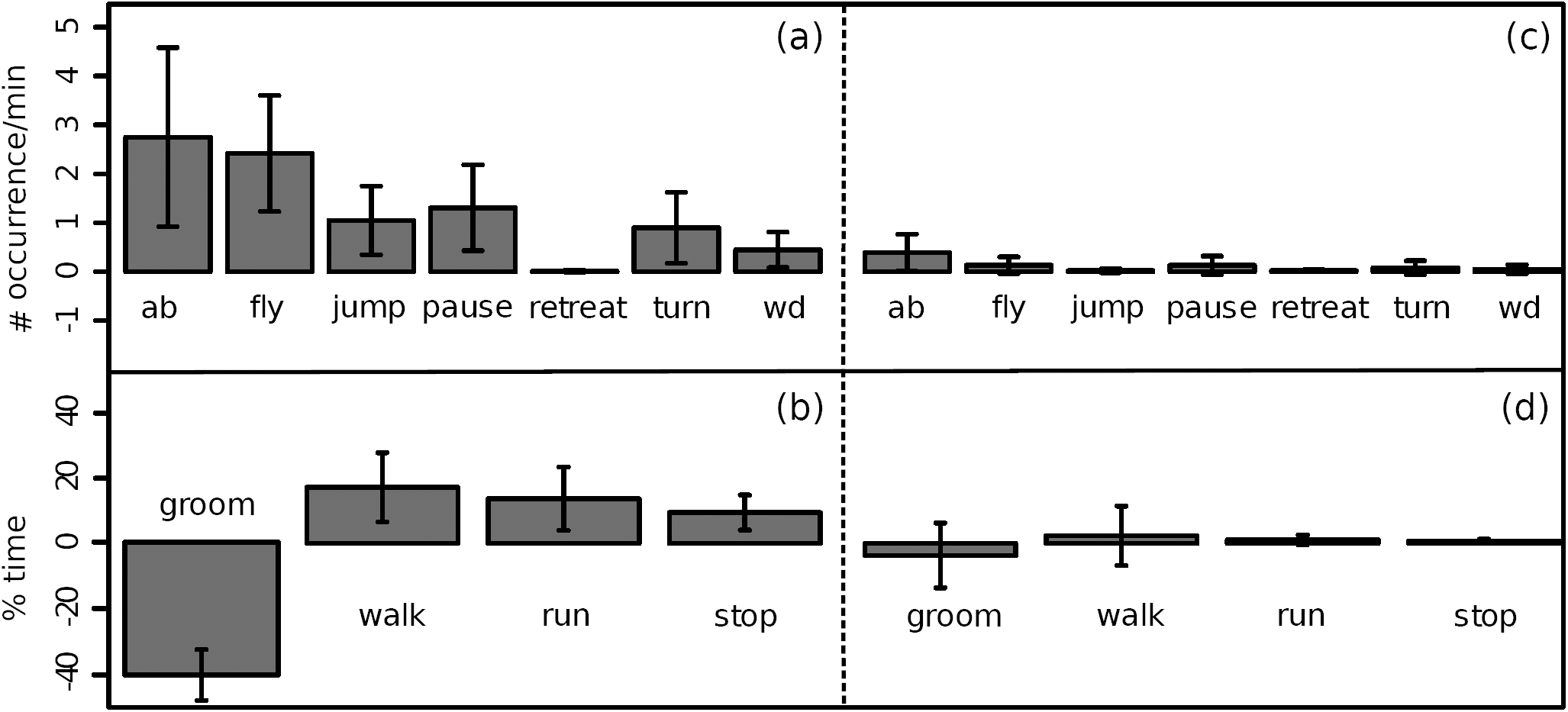
Fruit flies increase overall activity levels in the presence of jumping spiders (a) and (b) but not in the presence of mantids (c) and (d). Plots (a) and (b) show change in mean number of occurrences per minute of each behavioural state as a result of the addition of a predator. Plots (c) and (d) show mean change in percentage of total time spent in a given behavioural state caused by the addition of a predator. On the left of the dotted line, behavioural changes correspond to the presence of a spider whereas on the right of the dotted line, behavioural differences are due to the presence of a juvenile praying mantid. Error bars are ± 95% CI.

Given the design of our experiment, we were able to model the degree to which individuals varied in their responses to the jumping spiders. Individuals varied greatly both in their baseline activity levels as well as in their propensities to respond to jumping spiders. The among-individual coefficient of variation for time spent grooming in the absence of predators was 57.7% (40.1-74.2%). While most individuals reduced their grooming activity in the presence of predators, the degree to which they did so varied substantially, with the among individual coefficient of variation for the decrease being % (26.5-94.6%), as shown in Figure 3a. For walking, the among-individual coefficient of variation was 80.3% (50-105%) in the absence of the spider, and 135% (1-181%) for the magnitude of increase in the presence of the spider (Figure 3b).

**Figure 3.**
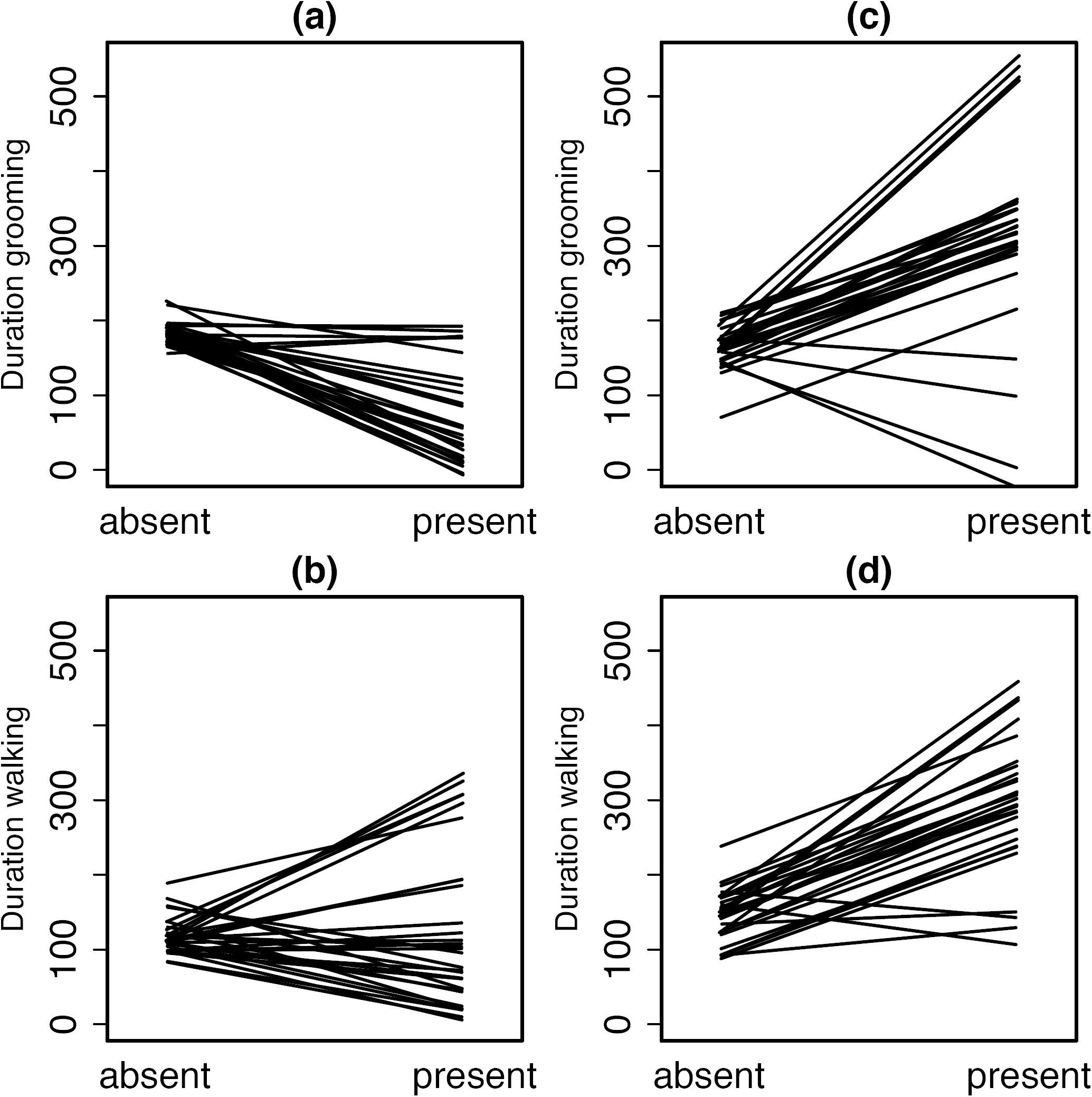
Inter individual behavioural variation in response to predators is present in natural populations. Reaction norms visualize how each individual fruit fly responded to the introduction of a spider (panels a and b) or a mantid (panels c and d) into the assay chamber. Measures are in seconds. Each line corresponds to response of one individual. Estimates are derived from the predicted values for each individual from the mixed models.

Performance of the stopping behaviour by *D. melanogaster* in the presence of spiders varied substantially among individuals, with the among-individual coefficient of variation being 168% (95% CI: 123-214%). This is driven in part by the fact that 40% of individual flies never performed stopping, even in the presence of the spider. There was a negative correlation (-0.84), between the amount of time individuals spent grooming before and after the addition of the spiders (Table 2). That is, on average, individuals who were more active prior to the addition of the spider reduced their activity to a greater extent in the presence of the spider. A similar negative correlation (-0.66) for among individual activity for locomotion, was observed (Table 2).

**Table 2.** Individual flies show negative correlations between behavioural states before and after the introduction of a predator. There is considerable variation among individuals in time spent performing specific behaviours (i.e. walking and grooming), with and without predators. However, there is a strong negative correlation within individuals for time spent before and after introduction of the predator. That is, individuals who spend more time performing a specific behaviour prior to the addition of a predator, reduce that behaviour to an even greater amount (than the average for the sample) once the predator is introduced. The one exception is for grooming for the mantid trials. Diagonals of the table contain the standard devation (mean of the posterior distribution) for individual behavioural responses (95% CIs in paratheses) from the random effects of the models. Above the diagonal are covariances between predictors (and CIs in parantheses). Below the diagonal are correlation coefficients for the covariances between the predictors.

**Grooming, Spider**
|  | **Intercept** | **Pred.State** | **Time** |
| --- | --- | --- | --- |
| **Intercept** | 89.2 (62.0, 115.3) | -68.2 (-88.7, -34.1) | 26.4 (14.0, 36.0) |
| **Pred.State** | -0.84 | 61.7 (18.8, 84.5 ) | -14 (-25.5, 17.1) |
| **Time** | 0.75 | -0.3 | 10.4 (4.4, 14.9 ) |

**Walking, Spider**
|  | **Intercept** | **Pred.State** | **Time** |
| --- | --- | --- | --- |
| **Intercept** | 81.9 (43.3, 109.8) | -60.2 (-89.5, 27.5) | 25.6 (-8.8, 36.7) |
| **Pred.State** | -0.66 | 67.3 (0.36, 98.5) | 11.5 (-22.2, 29.2) |
| **Time** | 0.62 | 0.15 | 13 (5.6, 18.2) |

**Grooming, Mantid**
|  | **Intercept** | **Pred.State** | **Time** |
| --- | --- | --- | --- |
| **Intercept** | 122.8 (59.6, 175) | -20.2 (-117, 106) | 46.0 (-16.9, 74.5) |
| **Pred.State** | -0.05 | 60.5 (0.13, 100.2) | -18.3 (-47.9, 34.0) |
| **Time** | 0.8 | -0.26 | 21.5 (2.6, 33.8) |

**Walking, Mantid**
|  | **Intercept** | **Pred.State** | **Time** |
| --- | --- | --- | --- |
| **Intercept** | 144.8 (86.2, 198) | -100.3 (-162.6, 38.8) | 63.2 (31.6, 90.3) |
| **Pred.State** | -0.86 | 80.5 (0.21, 139.3) | -45.2 (-76.4, 19.6) |
| **Time** | 0.94 | -0.86 | 29.4 (11.3, 43.5) |

To visualise the temporal associations among behavioural sequences, we constructed transition matrices (Supplement b Tables S1, S2, S5 and S6) and transition probability diagrams for all pairs of behaviours in the presence (Figure 4a) and absence (Supplement b Figure S7) of spiders. In response to jumping spiders, transitions among behaviours are somewhat more dispersed (with many connections between behaviours), suggesting that there is weak temporal association between fruit fly behaviours. Indeed these qualitative conclusions are supported based on the Z-scores. In the absence of spiders 8 possible transitions were significant (after controlling for multiple comparison, Supplement b Table S2), while 13 transitions were significant in the presence of the spider (Supplement b Table S1). Most of these differences were due to the increase in behaviours potentially involved with anti-predation activity (i.e. flight, abdominal lifting). However, while the results of the log-linear analysis (across the whole transition frequency matrix) supported the dependence of current behavioural states on the previous state (resid df=71, deviance=632, p < 0.001), the inclusion of predator status did not influence this dependence (resid df = 71, deviance = 59, p = 0.8).

**Figure 4.**
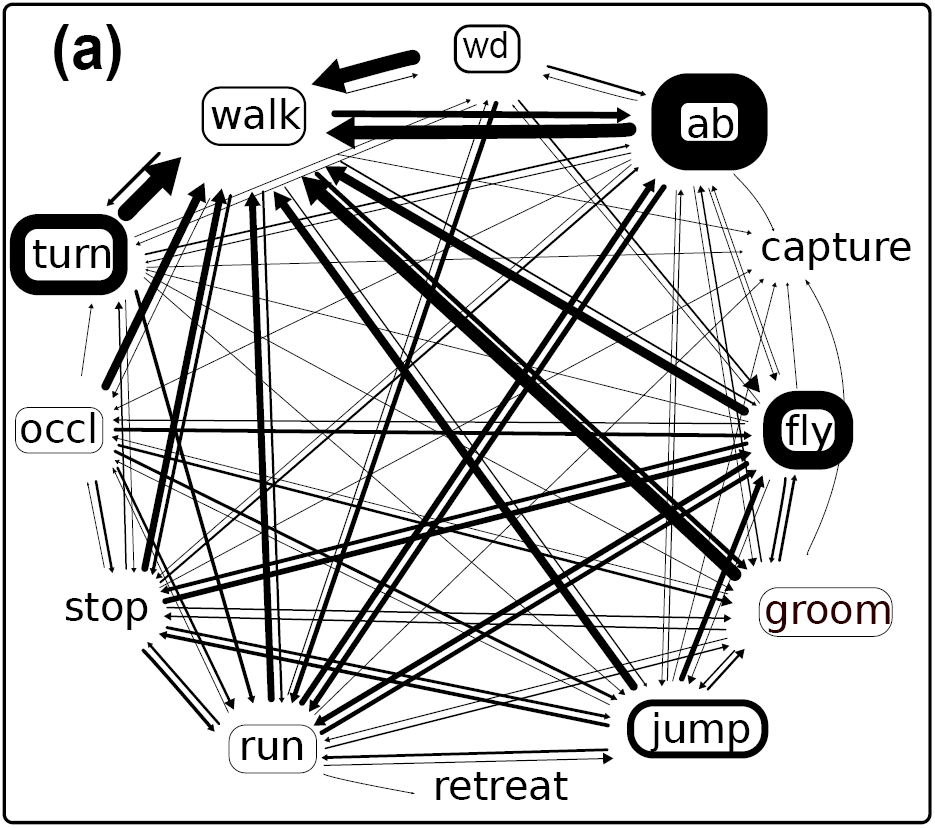
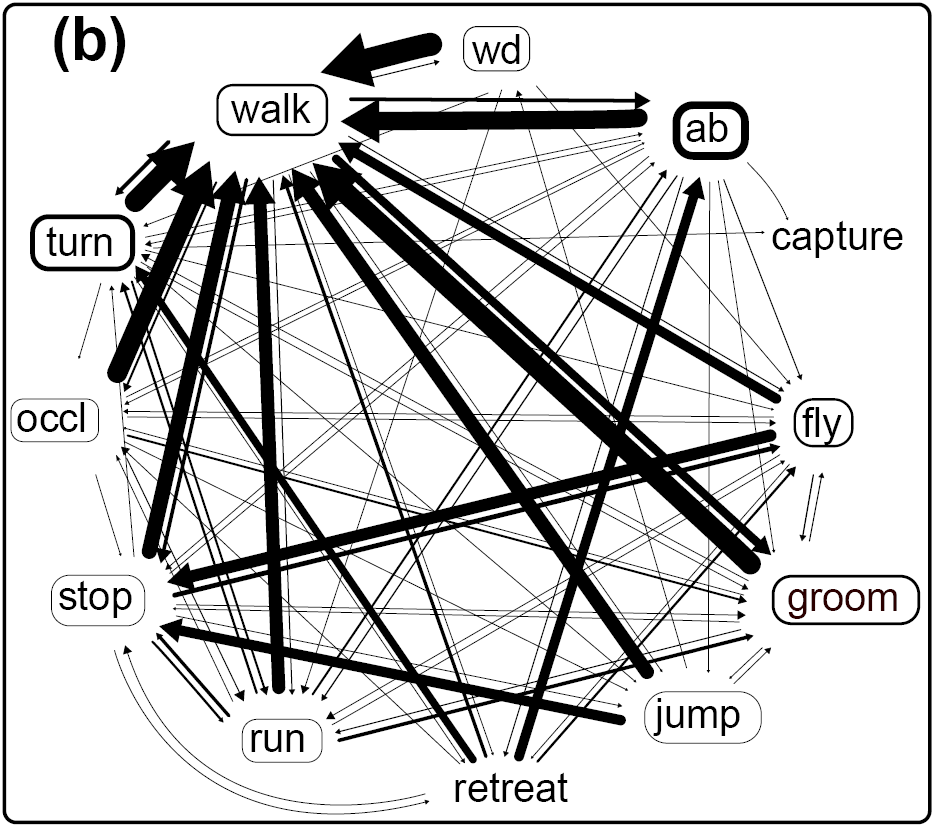
Spiders and mantids had different effects on the temporal associations between pairs of *D. melanogaster* behaviours. a) A diagram representing probability of transitioning from one fly behavioural state to the other in the presence of a zebra jumping spider. b) A diagram representing probability of transitioning from one fly behavioural state to the other in the presence of a juvenile praying mantid. Thickness of arrows indicates transition probability between the two behaviours. The arrowhead points to the behaviour being transitioned to. Thickness of the box around behavioural state (groom, run, occl, retreat, stop and walk) indicate the mean proportion of total time spent in that behaviour, whereas thickness of the box around behavioural events (fly, jump, turn, wd, ab) indicates mean number of occurrences per minute of that behaviour. To reduce the complexity of the web we combined the behaviours “pause” with the behaviour “stop”. Behavioural transitions that occurred less than 10 times have not been shown in the figure.

### Flies perform a previously undescribed retreat behaviour in response to mantids

In contrast to their behaviour in the presence of jumping spiders, the presence of a juvenile mantid had a minimal influence on *D. melanogaster*’s locomotory activity (Figure 2c and 2d, also see ethograms in Figure 1b and Supplement a). Time spent grooming, walking, running and stopping was highly variable (with estimates including zero) in the presence of a juvenile praying mantid (Figure 2d, Supplement b Figures S1 and S2). Grooming decreased by 20% (95% CI: 83% decrease to 36% increase), locomotion increased by 9% (13% decrease to 32% increase) and stopping increased by 40% (8% decrease to 85% increase). Similarly, the presence of a mantid did not influence the frequency at which *D. melanogaster* tended to perform most instantaneous behaviours (Figure 2c, Supplement b Figure S4). However, as was observed in the presence of spiders, flies substantially increased the frequency of abdominal lifting, from 0.45/minute to 5.6/min (95% CI: 2.56 - 12.17) in the presence of a juvenile mantid (Supplement b Figure S4). Upon encounter with a mantid, half of the individuals (15/30) performed a previously undescribed reversal behaviour (Supplement b video 3), which we term “retreat”. As with the zebra spiders, we saw no significant sex specific differences in response to mantids.

Although the presence of a mantid had a small effect on fly behaviour, flies did vary considerably in their grooming and walking activities. Indeed, the among-individual variability in proportion of time spent grooming and walking is greater in magnitude in the presence of the mantids than spiders (Figure 3). Evidence for negative co-variation for intra-individual behaviour before and after the addition of the predator was not strongly supported (i.e. 95% CIs for covariances included zero) (Table 2).

Transition matrices and transition probability diagrams (Supplement b Figure S7b, Figure 4b and Tables S3, S4, S7 and S8) show patterns of temporal association among behaviours. In response to juvenile mantids, the transitions diagram is less dispersed than that in the presence of jumping spiders (Figure 4), suggesting that the degree of association between behaviours in the presence of mantids is more extreme. While most behaviours (abdominal lift, fly, groom, jump, run, stop, and turn) tend to transition to walking, we also see stronger associations between other pairs of behaviours. For example, after performing the retreat behaviour, flies often either performed the abdominal lift or turn, while flight is often followed by stopping. These observations are supported by the findings that in the absence of mantids, 12 transitions showed significant deviations from expectations (Supplement b, Table S4). In comparison, in the presence of mantids 23 transitions showed a significant deviation from expected values (Supplement b, Table S3). Interestingly, as with the spiders the log-linear model supports the non-independence of behavioural states (resid df=71, deviance=1054, p <0.001), but not for the additional influence of predator state on this non-independence (resid df = 71, deviance = 72, p=0.4).

## DISCUSSION

Prey organisms can alter their behaviour to reduce the likelihood of detection, capture or encounter with a predator (Lima, 1998). For example, when predators are present, ground squirrels dedicate more time to vigilance behaviours (like scanning for a predator, see (Bachman, 1993) and some aquatic insects spend more time in refuges (Kohler & McPeek, 1989). These changes in behaviour may alter the use of resources, and potentially the fitness of an organism. However, the nature and intensity of non-consumptive effects of a predator on its prey are a function of several predator specific factors, one of which is the predator’s hunting mode (Preisser, Orrock & Schmitz, 2007). Predator hunting mode, i.e., the set of behavioural strategies that a predator employs to pursue and capture its prey (Schoener, 1971; Huey & Pianka, 1981; Schmitz, 2008) can be an important determinant of a prey organism’s anti-predatory behavioural response (Schmitz, 2008). In this study, we describe the anti-predatory behavioural repertoire of a natural population of *Drosophila melanogaster* in response to predation by the zebra jumping spider (*Salticus scenicus)* and juvenile Chinese praying mantids (*Tenodera aridifolia sinensis*). Among other characteristics, zebra spiders and praying mantids differ in their hunting mode. While we discuss our findings with respect to hunting mode differences, we recognize that other attributes differing among the predators may contribute to the observed differences in prey behavioural repertoires. However, as our experimental design was meant to minimize the effects of many possible confounding factors (e.g. time of day, temperature, humidity) it seems likely that, in part, our results reflect hunting mode differences.

In response to active hunters (those that constantly patrol for prey), we predicted that fruit flies would increase their overall activity levels (including flight) in order to maintain maximum distance from the predator at all times; To reduce the likelihood of an encounter with an ambush predator however (i.e., a predator that only attacks when a prey organism wanders in to its strike zone), we predicted that *D. melanogaster* would respond by decreasing locomotory activities (Schmitz, 2008). Our results, however, were only partially in line with these predictions. While the actively hunting jumping spiders induce a clear increase in overall activity, we found the presence of juvenile mantids-our ambush predators-to have minimal influence on fruit fly activity levels (Figure 2, Supplement b Figure S2). It has been previously argued that ambush predators might be a predictable source of threat to prey organisms (Preisser, Orrock & Schmitz, 2007; Schmitz, 2008) as opposed to the diffuse and variable threat imposed by active hunters (Schmitz, 2008). Therefore, it is perhaps surprising that fruit flies show a stronger behavioural response to the threat of active hunters (zebra jumping spiders). However, our predictions are based on studies on a grasshopper and its two predatory spider species that differ in hunting mode. Given that selection pressures faced by adult diptera are different from those experienced by grasshoppers (orthoptera), such predictions may not be generalizable. Several factors including body size and dispersal patterns may contribute to this difference. Many species of jumping spiders, including *S. scenicus,* are often seen in the natural habitat of *D. melanogaster* (personal observations of A.P., C.P. and I.D.), and are likely to be ecologically relevant predators of *Drosophila*. Mantids however, are rarely found in areas where fruit flies are abundant (personal observations of A.P. and I.D.), at least in Eastern North America. Therefore, it is likely that fruit flies, having experienced a longer evolutionary history with small jumping spiders, are better able to recognize these spiders as a threat. In addition, the disturbance created by a constantly patrolling zebra spider may be partly responsible for the increased activity levels seen in *D. melanogaster* (either due to actual mechanical disturbance or because flies are able to detect moving objects quicker than stationary ones). In this study, we are unable to tease apart the effects of evolutionary recognition versus constant mechanical disturbance on the differences in flies’ activity levels. Further experimentation with harmless but constantly moving heterospecifics (such as field crickets) or immobilized active hunters might be useful in addressing these issues.

We also identified a number of (to our knowledge) undescribed behaviours of *D. melanogaster,* potentially relating to its interactions with predators. The behaviour we called “stopping” (Table 1) was observed numerous times after a direct (but failed) attack by a spider (Supplement 3 video 1). While *D. melanogaster* will spend time without any ambulatory activity (walking, running), they are almost always observed to be active (generally grooming) during these periods. However, when fruit flies performed the stopping behaviour, there was a complete lack of movement on the part of the fly, even when video was viewed at a few frames/second. When a fruit fly was “stopped”, the spider had to search for the fly, irrespective of the physical proximity between the spider and the fly. In salticids, while the principal eyes have high spatial acuity, secondary eyes are primarily used to detect moving objects (Harland, Jackson & Macnab, 1999; Land, 1971). Because salticids are unable to accommodate by changing the shape of their lens, they need to extensively sample their visual field to see details in object shape and form (Harland, Jackson & Macnab, 1999; Land, 1971; Blest, Hardie & McIntyre, 1981). Scanning for prey by such sampling is likely a slow process unless guided by the motion sensing peripheral eyes, giving motionless prey the advantage of staying hidden (at least for a few seconds) while in plain sight of their salticid predator. Thus, *D. melanogaster* may be using the “stopping” behaviour as a potential mechanism to reduce the likelihood of detection by the spider.

Additionally, in the presence of both predators, *D. melanogaster* substantially increase the frequency at which it performed abdominal lifts. To our knowledge, abdominal lifting has not been described in *D. melanogaster* literature before and may be relevant in an anti-predatory context. While studying courtship behaviours in female *D. melanogaster*, Lasbleiz, Ferveur & Everaerts (2006) described two behaviours perhaps similar to the abdominal lifting described here: abdominal drumming and abdominal extension. Abdominal drumming (described as “quickly repeated vertical movements of the abdomen which is tapped on the substrate”) was only seen in males during courtship display, and abdominal extensions (described as “abdomen raised by 15-30 degrees”) were also closely associated with courtship. Because abdominal lifting was often directed at a predator or followed a failed predatory encounter, we suspect abdominal lifting to be different from abdominal extensions and abdominal drumming, and with a possibly anti-predatory function. We speculate that if abdominal lifting is indeed anti-predatory, it could function in one of several possible ways. First, abdominal lifting may be a signal of prey condition directed at the predator as a form of pursuit deterrence, comparable to stotting in the Thomson’s gazelle (FitzGibbon & Fanshawe, 1988). Second, because *D. melanogaster* are often surrounded by conspecifics, abdominal lifting may be a means though which one fly warns its conspecifics of the presence of a potential threat (similar in function to fin flicking in tetras, (Brown, Godin & Pedersen, 1999). Finally it may be an indication of some sort of physiological priming of the fly in preparation for a fight-or-flight response. Determining whether it is a specific anti-predator behaviour, as well as the details of its function need to be a focus of future work.

In response to the juvenile praying mantids, half of the fruit flies we observed (15/30) performed a reverse walking behaviour which we have called “retreat”, where the flies walked in reverse, away from the predator (supplement b, video 3). This was often (but not always) interspersed with the abdominal lifting behaviour. Phenomenologically, this behaviour may be similar to that described in Bidaye et al. (2014). Bidaye el al. (2014) identified neurons that upon activation changed walking direction in *D. melanogaster*. Bidaye et al’s reverse walking behaviour appears to be a smooth and continuous behaviour, whereas “retreat” was often discontinuous and accompanied by abdominal lifting. If the two “retreat” behaviours are related, the observed disassociation between retreat and abdominal lifting as well as its continuous nature (in Bidaye el al., 2014) may be a function of how the neurons were perturbed.

We also investigated how the presence of the different predators may influence non-random associations among behaviours. We observed that in the presence of both predators there was an increase in the number of behavioural transitions that deviated from expectations under independence (from 12 to 23 with the mantid, and 8 to 13 with the spider). Despite this, the log-linear model (analysing the whole transition frequency matrix) did not support the influence of predator state on the frequencies of transitions. This may be partly due to the relatively modest sample sizes (in terms of both individuals and transitions among behaviours). Further work is necessary to validate and extend this sequential analysis.

While we show that there are some predator hunting-mode specific behavioural differences in *D. melanogaster’s* anti-predator response, we reiterate two important caveats. First, although the primary distinction between the zebra jumping spider and juvenile Chinese praying mantids as predators is their hunting-mode, other factors between these species (for example, size, colour, odour) may also influence differences in fruit fly behaviours. Replicating the observations with other predator pairs that differ in hunting-mode is necessary to confirm hunting-mode’s influence on anti-predatory repertoires. Secondly, our assay chambers are an artificial environment and do not resemble the conditions under which *D. melanogaster* face predators in the wild. Due to the nature of our assay chamber, *D. melanogaster* were unable to employ behavioural strategies that may reduce encounters with predators (e.g., utilizing a refuge). Therefore we were only able to describe the capture-deterrence repertoire of *D. melanogaster* behaviour. We believe that our study is a necessary first step to describing and documenting the complete anti-predatory behavioural repertoire of *D. melanogaster* and we foresee future work to be conducted in a modified chamber, under more “natural” conditions. Doing so will allow us to take this premier model genetic system and make it into an ecological model as well.

## ACKNOWLEDGEMENTS

We thank Dr. Fred Dyer for loaning us video cameras. Thanks to Dr. Ben DeBivort for discussions and suggestions and to Dr. Reuven Dukas and Dr. Marla Sokolowski and members of the Dworkin lab for comments on the manuscript. This material is based in part upon work supported by the National Science Foundation under Cooperative Agreement No. DBI-0939454. Any opinions, findings, and conclusions or recommendations expressed in this material are those of the author(s) and do not necessarily reflect the views of the National Science Foundation.

## Supplemental material

### Table of contents

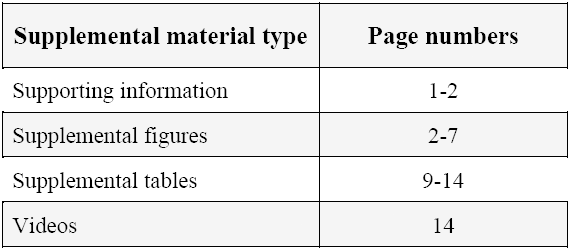

### Supporting information

#### Differential response to spiders versus mantids

Because spider and mantid population densities vary by season, we had to temporally segregate the spider assays from the mantid assays. We conducted all spider observations between October and December 2012 and all the mantid observations between March and May 2013. Comparing time allocation and frequencies of occurrences in the predator absent state between the two predator treatments suggest that behavioural modifications were predator induced, and not due to seasonal effects (Figure S5 and S6). Although the assays were carried out under highly controlled conditions, to confirm that predator species-specific behavioural differences were not confounded with seasonal differences in behaviour, we performed 6 additional assays (alternating between spider and mantid treatments) within the span of one week. The control experiments show no evidence of confounding effects of season with *D. melanogaster*’s anti-predator behavioural repertoire (Table S9, S11 and S12 below). Ethograms are shown in Supplement a. Furthermore, to confirm that the disturbance we caused (to the assay chamber) during the addition of a predator did not confound behavioural responses to the predator, we did 3 “no predator” control assays. For these “no predator” controls, instead of adding a predator to the arena, we caused a mild disturbance (∼ to intensity of disturbance caused while adding the predator) without actually adding any predator. We found that disturbance caused during predator addition was not responsible for observed behavioural modifications (Table S10 and S13). Finally, “no predator” controls also ruled our temporal differences in fruit fly activity levels (Table S10 and S13)

### Supplemental figures

**Figure S1.**
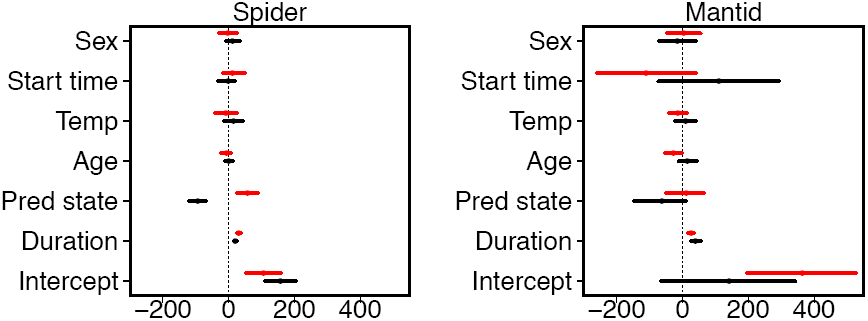
Presence of jumping spiders causes fruit flies to walk more and groom less, whereas the presence of mantids causes weaker, more variable (and not significant) changes in fruit fly activity levels. Here we show coefficient plots from the output of mixed effects models using the package MCMCglmm to visualize duration of two behaviours (**grooming in black** and **locomotion in red**) as a function of predator state (present vs absent of spiders, left panels and mantids, right panels), time spent in the assay [Time (cent.)] total, sex of the fly, start time of the assay, temperature in the room and age of the fly. The continuous covariate “Time” was centered around the mean and therefore reflects the average increase in time spent per minute of the assay. Estimates are in seconds. Error bars are ± 95% CI.

**Figure S2.**
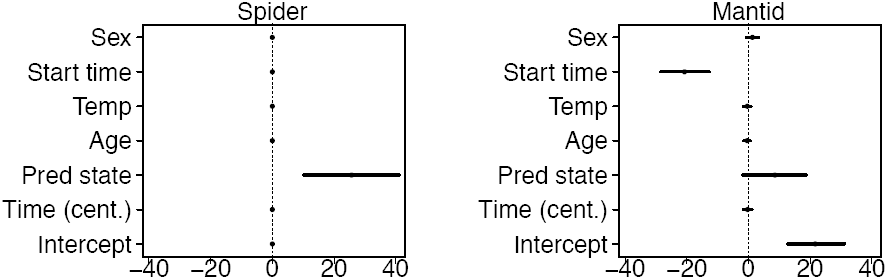
Fruit flies “stop” significantly longer in the presence of spiders (left panel), and to a much lesser extent (and not significantly) in the presence of mantids (right panel). Here we have coefficient plots made from the output of mixed effects models using the package MCMCglmm to visualize duration of “stopping” as a function of predator state (present vs absent), time spent in the assay [Time (cent.)], sex of the fly, start time of the assay, temperature in the room and age of the fly. Estimates are in seconds. Error bars are ± 95% CI. The continuous covariate “Time” was centered around the mean and therefore reflects the average increase in time spent per minute of the assay. Although assays were performed between 9 am and 12 pm each day, start time for the mantid assays significantly affected the total time that flies spent “stopping”.

**Figure S3.**
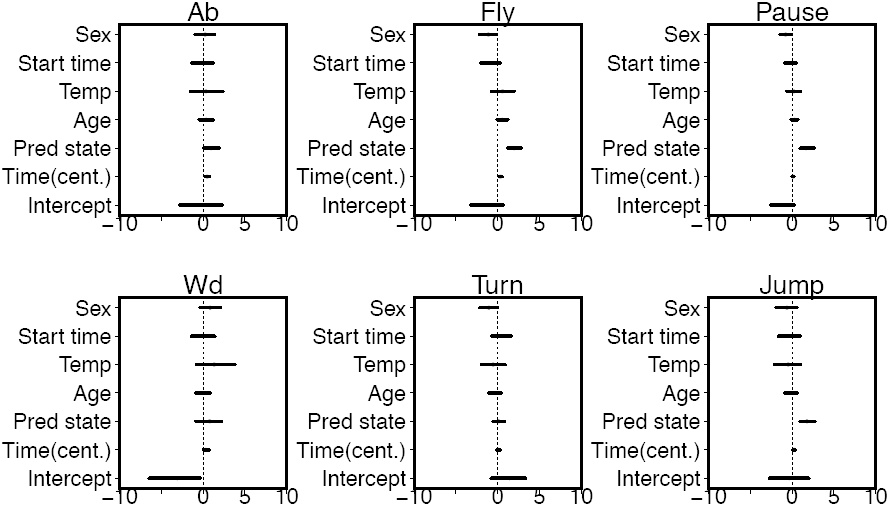
In the presence of spiders, fruit flies increased the frequency with which they performed flights, pauses and jumps. Here we show coefficient plots made from the output of mixed effects models where the events were modeled using a poisson generalized linear mixed model with a log-link function fit using the `MCMCglmm` function, and estimates remain on a natural log scale. Coefficient plots were used to visualize frequency of each individual behavioural event (ab, fly, pause, wd, turn and jump) as a function of predator state (present vs absent of a spider), time spent in the assay [Time (cent.)], sex of the fly, start time of the assay, temperature in the room and age of the fly. All estimates are scaled to number of events per minute. Error bars are ± 95% CI.

**Figure S4.**
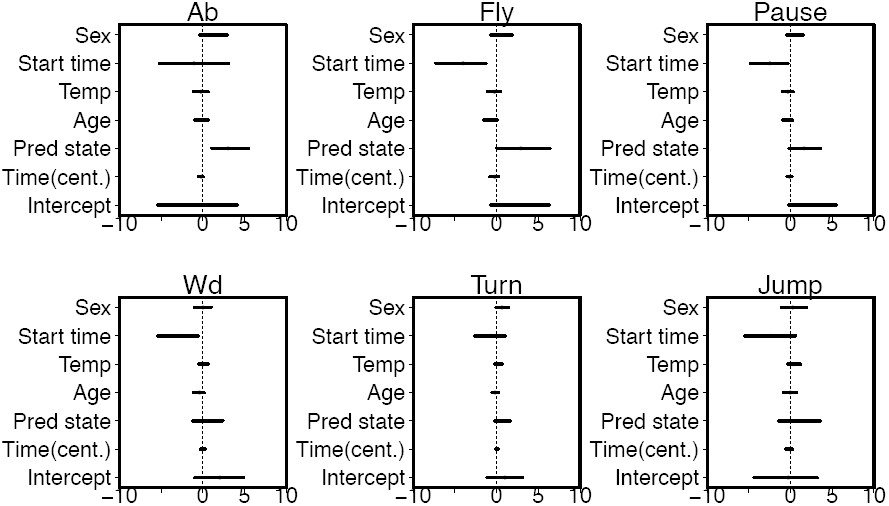
Fruit flies performed abdominal lifts are a higher frequency in the presence of a juvenile mantid. Coefficient plots made from the output of mixed effects models where the events were modeled using a poisson generalized linear mixed model with a log-link function fit using the MCMCglmm function, and estimates remain on a natural log scale. Coefficient of the model output are used to visualize frequency of each individual behavioural event (ab, fly, pause, wd, turn and jump) as a function of predator state (present vs absent of a mantid), time spent in the assay [Time (cent.)], sex of the fly, start time of the assay, temperature in the room and age of the fly. All estimates are scaled to number of events per minute. Error bars are ± 95% CI. Although assays were performed between 9 am and 12 pm each day, start time for the mantid assays significantly affected the frequency at which *D. melanogaster* performed the “Fly”, “Wd” and “Jump” behaviours.

**Figure S5.**
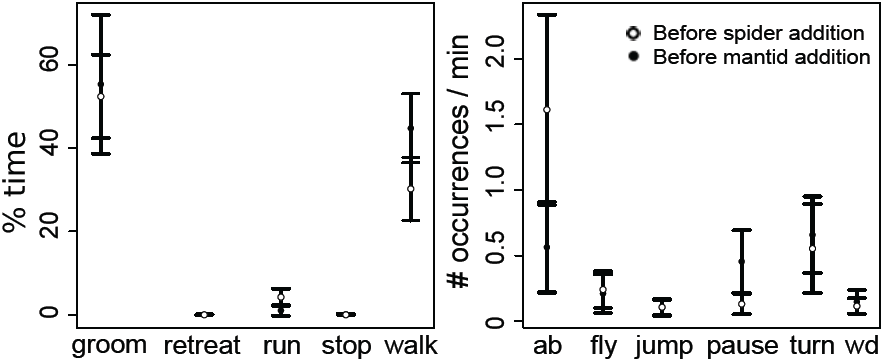
Hunting mode induced behavioural differences in fruit fly behaviours were not confounded with seasonal effects. Here we show percentage time spent in each behavioural state (left) and number of occurrences per minute for each behavioural event (right) as measured for individual fruit flies before the addition of a spider (white circles) and before the introduction of a mantid (black circles) into the chamber. Error bars are ± 2 * SEs. Overlapping error bars suggest that there was minimal effect of season on the behavioural repertoire of fruit flies.

**Figure S6.**
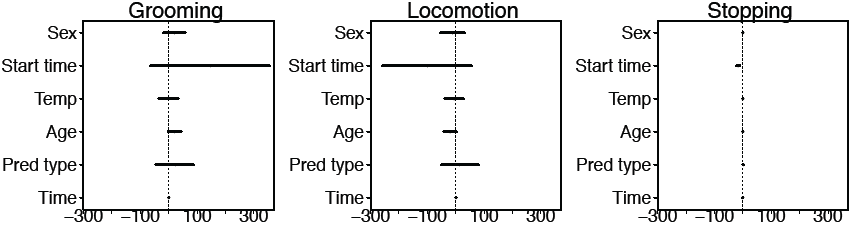
Seasonal differnces in fruit fly behaviours did not confound behavioural differences induced by different hunting-modes. Flies measured before the addition of a spider did not differ in behaviour from flies measured before the addition of a mantid.

**Figure S7.**
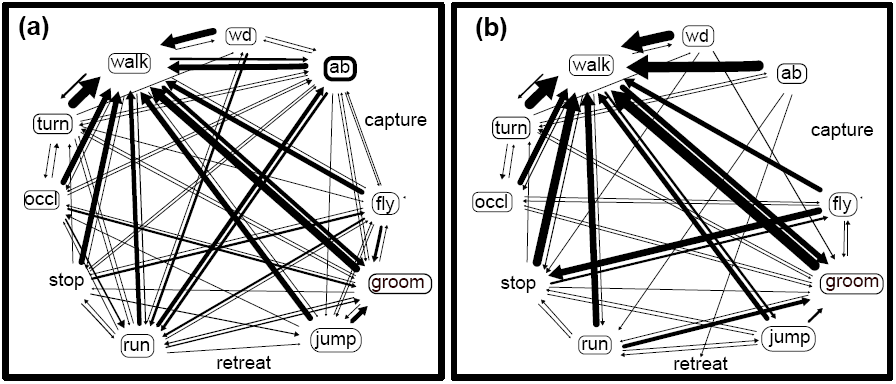
**a)** A diagram representing probability of transitioning from one fly behaviour to the other when individuals were measured before the addition of a spider **b)** A diagram representing probability of transitioning from one fly behaviour to the other for individuals measured before the addition of a juvenile mantid. Thickness of arrows indicates transition probability between the two behaviours. The arrowhead points to the behaviour being transitioned to. Thickness of the box around behavioural state (groom, run, occl, retreat, stop and walk) indicate the mean proportion of total time spent in that behaviour, whereas thickness of the box around behavioural events (fly, jump, turn, wd, ab) indicates mean number of occurrences per minute of that behaviour. To reduce the complexity of the web we combined the behaviours “pause” with the behaviour “stop”. Behavioural transitions that occurred less than 10 times have not been shown in the figure.

### Supplemental tables

**Table S1.** Transition frequency matrix when a spider was present in the chamber. Each row represents the number of times one behaviour (row name) transitioned to another behaviour (column name). Numbers in blue represent transitions that occurred more often that expected under a model of independence, whereas numbers in red are transitions that occurred less often than expected (see methods).

| Spider |  |  |  |  |  |  |  |  |  |  |  |  |  |
| --- | --- | --- | --- | --- | --- | --- | --- | --- | --- | --- | --- | --- | --- |
| Behavior | *ab* | *capture* | *fly* | *groom* | *jump* | *retreat* | *run* | *stop* | *occl* | *turn* | *walk* | *wd* | Total |
| *ab* | 0 | 2 | 8 | 9 | 7 | 0 | 45 | 6 | 5 | 10 | 112 | 7 | 211 |
| *capture* | 0 | 0 | 0 | 0 | 0 | 0 | 0 | 0 | 0 | 0 | 0 | 0 | 0 |
| *fly* | 4 | 2 | 0 | 15 | 7 | 0 | 34 | 25 | 10 | 1 | 58 | 1 | 157 |
| *groom* | 11 | 3 | 16 | 0 | 10 | 0 | 8 | 11 | 5 | 7 | 77 | 2 | 150 |
| *jump* | 4 | 2 | 19 | 8 | 0 | 0 | 9 | 13 | 2 | 0 | 28 | 0 | 85 |
| *retreat* | 1 | 0 | 0 | 0 | 0 | 0 | 0 | 0 | 0 | 0 | 0 | 0 | 1 |
| *run* | 43 | 4 | 33 | 13 | 9 | 1 | 0 | 18 | 3 | 4 | 51 | 9 | 188 |
| *stop* | 9 | 3 | 32 | 10 | 16 | 0 | 19 | 0 | 1 | 9 | 38 | 0 | 137 |
| *occl* | 3 | 0 | 6 | 4 | 5 | 0 | 4 | 4 | 0 | 1 | 14 | 0 | 41 |
| *turn* | 8 | 1 | 0 | 4 | 3 | 0 | 14 | 2 | 0 | 0 | 66 | 2 | 100 |
| *walk* | 122 | 7 | 36 | 87 | 26 | 0 | 42 | 60 | 14 | 63 | 0 | 24 | 481 |
| *wd* | 5 | 0 | 4 | 0 | 0 | 0 | 9 | 0 | 0 | 1 | 29 | 0 | 48 |
| Total | 210 | 24 | 154 | 150 | 83 | 1 | 184 | 139 | 40 | 96 | 473 | 45 | 1599 |

**Table S2.** Transition frequency matrix before a spider was added to the chamber. Each row represents the number of times one behaviour (row name) transitioned to another behaviour (column name). Numbers in blue represent transitions that occurred more often that expected under a model of independence, whereas numbers in red are transitions that occurred less often than expected (see methods)s.

No spider
| Behavior | ab | capture | fly | groom | jump | retreat | run | stop | occl | turn | walk | wd | Total |
| --- | --- | --- | --- | --- | --- | --- | --- | --- | --- | --- | --- | --- | --- |
| ab | 0 | 0 | 2 | 16 | 1 | 0 | 30 | 1 | 2 | 10 | 65 | 2 | 129 |
| capture | 0 | 0 | 0 | 0 | 0 | 0 | 0 | 0 | 0 | 0 | 0 | 0 | 0 |
| fly | 1 | 0 | 0 | 9 | 1 | 0 | 6 | 2 | 0 | 0 | 15 | 0 | 34 |
| groom | 7 | 0 | 11 | 0 | 8 | 0 | 18 | 1 | 5 | 10 | 122 | 0 | 182 |
| jump | 0 | 0 | 1 | 7 | 0 | 0 | 0 | 0 | 1 | 0 | 9 | 0 | 18 |
| retreat | 0 | 0 | 0 | 0 | 0 | 0 | 0 | 0 | 0 | 0 | 0 | 0 | 0 |
| run | 38 | 0 | 5 | 14 | 1 | 0 | 0 | 4 | 2 | 1 | 42 | 3 | 110 |
| stop | 1 | 0 | 4 | 0 | 2 | 0 | 2 | 0 | 0 | 1 | 10 | 0 | 20 |
| occl | 2 | 0 | 0 | 6 | 0 | 0 | 5 | 0 | 0 | 3 | 16 | 0 | 32 |
| turn | 4 | 0 | 1 | 5 | 0 | 0 | 2 | 0 | 3 | 0 | 72 | 1 | 88 |
| walk | 74 | 0 | 8 | 126 | 5 | 0 | 42 | 13 | 18 | 64 | 0 | 12 | 362 |
| wd | 1 | 0 | 0 | 0 | 0 | 0 | 4 | 0 | 0 | 0 | 13 | 0 | 18 |
| Total | 128 | 0 | 32 | 183 | 18 | 0 | 109 | 21 | 31 | 89 | 364 | 18 | 993 |

**Table S3.** Transition frequency in the presence of a juvenile praying mantid. Each row represents the number of times one behaviour (row name) transitioned to another behaviour (column name). Numbers in blue represent transitions that occurred more often that expected whereas numbers in red are transitions that occurred less often than expected.

Mantid
| Behavior | ab | capture | fly | groom | jump | retreat | run | stop | occl | turn | walk | wd | Total |
| --- | --- | --- | --- | --- | --- | --- | --- | --- | --- | --- | --- | --- | --- |
| ab | 0 | 1 | 9 | 2 | 1 | 9 | 10 | 5 | 4 | 6 | 127 | 0 | 174 |
| capture | 0 | 0 | 0 | 0 | 0 | 0 | 0 | 0 | 0 | 0 | 0 | 0 | 0 |
| fly | 0 | 0 | 0 | 7 | 0 | 1 | 5 | 60 | 2 | 0 | 50 | 0 | 125 |
| groom | 0 | 0 | 17 | 0 | 3 | 0 | 18 | 3 | 5 | 13 | 305 | 1 | 365 |
| jump | 0 | 0 | 0 | 1 | 0 | 0 | 0 | 13 | 0 | 0 | 18 | 1 | 33 |
| retreat | 9 | 0 | 2 | 0 | 0 | 0 | 0 | 1 | 1 | 7 | 3 | 0 | 23 |
| run | 11 | 0 | 3 | 18 | 0 | 0 | 0 | 17 | 1 | 12 | 68 | 0 | 130 |
| stop | 4 | 0 | 45 | 9 | 4 | 5 | 21 | 0 | 0 | 5 | 139 | 0 | 232 |
| occl | 1 | 0 | 3 | 7 | 1 | 0 | 5 | 1 | 0 | 0 | 73 | 0 | 91 |
| turn | 4 | 1 | 2 | 6 | 1 | 3 | 15 | 0 | 1 | 0 | 164 | 0 | 197 |
| walk | 145 | 2 | 43 | 326 | 23 | 5 | 51 | 132 | 73 | 150 | 0 | 49 | 999 |
| wd | 0 | 0 | 1 | 0 | 0 | 0 | 2 | 0 | 0 | 1 | 47 | 0 | 51 |
| Total | 174 | 4 | 125 | 376 | 33 | 23 | 127 | 232 | 87 | 194 | 994 | 51 | 2420 |

**Table S4.** Transition frequency matrix before a juvenile mantid was added to the chamber. Each row represents the number of times one behaviour (row name) transitioned to another behaviour (column name). Numbers in blue represent transitions that occurred more often that expected whereas numbers in red are transitions that occurred less often than expected.

No mantid
| Behavior | ab | capture | fly | groom | jump | retreat | run | stop | occl | turn | walk | wd | Total |
| --- | --- | --- | --- | --- | --- | --- | --- | --- | --- | --- | --- | --- | --- |
| ab | 0 | 0 | 0 | 0 | 0 | 1 | 1 | 0 | 0 | 1 | 31 | 0 | 34 |
| capture | 0 | 0 | 0 | 0 | 0 | 0 | 0 | 0 | 0 | 0 | 0 | 0 | 0 |
| fly | 0 | 0 | 0 | 1 | 0 | 0 | 0 | 19 | 1 | 0 | 11 | 0 | 32 |
| groom | 0 | 0 | 9 | 0 | 1 | 0 | 4 | 1 | 3 | 12 | 212 | 0 | 242 |
| jump | 0 | 0 | 0 | 3 | 0 | 0 | 1 | 4 | 0 | 0 | 6 | 0 | 14 |
| retreat | 1 | 0 | 0 | 0 | 0 | 0 | 0 | 1 | 1 | 0 | 0 | 0 | 3 |
| run | 0 | 0 | 0 | 9 | 1 | 0 | 0 | 1 | 0 | 0 | 15 | 0 | 26 |
| stop | 0 | 0 | 14 | 1 | 1 | 0 | 1 | 0 | 0 | 1 | 54 | 0 | 72 |
| occl | 0 | 0 | 1 | 2 | 0 | 0 | 0 | 0 | 0 | 1 | 55 | 0 | 59 |
| turn | 2 | 0 | 0 | 2 | 0 | 0 | 0 | 0 | 4 | 0 | 79 | 0 | 87 |
| walk | 31 | 0 | 8 | 236 | 10 | 1 | 19 | 45 | 48 | 71 | 0 | 21 | 490 |
| wd | 0 | 0 | 0 | 2 | 0 | 0 | 0 | 1 | 0 | 0 | 18 | 0 | 21 |
| Total | 34 | 0 | 32 | 256 | 13 | 2 | 26 | 72 | 57 | 86 | 481 | 21 | 1080 |

**Table S5.** Transition probability from one behaviour (row name) to the other (column name) in the presence of a zebra jumping spider. Transition probabilites are obtained by dividing each transition frequeny (see table S1) between a pair of behaviours by the total number of times a given behaviour was performed (row sums in table S1).

Spider
| Behavior | ab | capture | fly | groom | jump | retreat | run | stop | occl | turn | walk | wd |
| --- | --- | --- | --- | --- | --- | --- | --- | --- | --- | --- | --- | --- |
| ab | 0.00 | 0.01 | 0.04 | 0.04 | 0.03 | 0.00 | 0.22 | 0.03 | 0.02 | 0.05 | 0.55 | 0.03 |
| capture | 0.00 | 0.00 | 0.00 | 0.00 | 0.00 | 0.00 | 0.00 | 0.00 | 0.00 | 0.00 | 0.00 | 0.00 |
| fly | 0.03 | 0.01 | 0.00 | 0.10 | 0.04 | 0.00 | 0.22 | 0.16 | 0.06 | 0.01 | 0.37 | 0.01 |
| groom | 0.07 | 0.02 | 0.11 | 0.00 | 0.07 | 0.00 | 0.05 | 0.07 | 0.03 | 0.05 | 0.52 | 0.01 |
| jump | 0.05 | 0.02 | 0.22 | 0.09 | 0.00 | 0.00 | 0.11 | 0.15 | 0.02 | 0.00 | 0.33 | 0.00 |
| retreat | 1.00 | 0.00 | 0.00 | 0.00 | 0.00 | 0.00 | 0.00 | 0.00 | 0.00 | 0.00 | 0.00 | 0.00 |
| run | 0.24 | 0.02 | 0.18 | 0.07 | 0.05 | 0.01 | 0.00 | 0.10 | 0.02 | 0.02 | 0.28 | 0.05 |
| stop | 0.07 | 0.02 | 0.23 | 0.07 | 0.12 | 0.00 | 0.14 | 0.00 | 0.01 | 0.07 | 0.28 | 0.00 |
| occl | 0.07 | 0.00 | 0.15 | 0.10 | 0.12 | 0.00 | 0.10 | 0.10 | 0.00 | 0.02 | 0.34 | 0.00 |
| turn | 0.08 | 0.01 | 0.00 | 0.04 | 0.03 | 0.00 | 0.14 | 0.02 | 0.00 | 0.00 | 0.67 | 0.02 |
| walk | 0.27 | 0.02 | 0.08 | 0.19 | 0.06 | 0.00 | 0.09 | 0.13 | 0.03 | 0.14 | 0.00 | 0.05 |
| wd | 0.10 | 0.00 | 0.08 | 0.00 | 0.00 | 0.00 | 0.19 | 0.00 | 0.00 | 0.02 | 0.60 | 0.00 |

**Table S6.** Transition probability from one behaviour (row name) to the other (column name) before a zebra jumping spider was introduced into the arena. Transition probabilites are obtained by dividing each transition frequeny (see table S1) between a pair of behaviours by the total number of times a given behaviour was performed (row sums in table S1).

| Behavior | *ab* | *capture* | *fly* | *groom* | *jump* | *retreat* | *run* | *stop* | *occl* | *turn* | *walk* | *wd* |
| --- | --- | --- | --- | --- | --- | --- | --- | --- | --- | --- | --- | --- |
| *ab* | 0.00 | 0.00 | 0.02 | 0.13 | 0.01 | 0.00 | 0.24 | 0.01 | 0.02 | 0.08 | 0.51 | 0.02 |
| *capture* | 0.00 | 0.00 | 0.00 | 0.00 | 0.00 | 0.00 | 0.00 | 0.00 | 0.00 | 0.00 | 0.00 | 0.00 |
| *fly* | 0.03 | 0.00 | 0.00 | 0.26 | 0.03 | 0.00 | 0.18 | 0.06 | 0.00 | 0.00 | 0.44 | 0.00 |
| *groom* | 0.04 | 0.00 | 0.06 | 0.00 | 0.04 | 0.00 | 0.10 | 0.01 | 0.03 | 0.05 | 0.67 | 0.00 |
| *jump* | 0.00 | 0.00 | 0.06 | 0.39 | 0.00 | 0.00 | 0.00 | 0.00 | 0.06 | 0.00 | 0.50 | 0.00 |
| *retreat* | 0.00 | 0.00 | 0.00 | 0.00 | 0.00 | 0.00 | 0.00 | 0.00 | 0.00 | 0.00 | 0.00 | 0.00 |
| *run* | 0.36 | 0.00 | 0.05 | 0.13 | 0.01 | 0.00 | 0.00 | 0.04 | 0.02 | 0.01 | 0.39 | 0.03 |
| *stop* | 0.05 | 0.00 | 0.20 | 0.00 | 0.10 | 0.00 | 0.10 | 0.00 | 0.00 | 0.05 | 0.50 | 0.00 |
| *occl* | 0.06 | 0.00 | 0.00 | 0.19 | 0.00 | 0.00 | 0.16 | 0.00 | 0.00 | 0.09 | 0.50 | 0.00 |
| *turn* | 0.05 | 0.00 | 0.01 | 0.06 | 0.00 | 0.00 | 0.02 | 0.00 | 0.03 | 0.00 | 0.83 | 0.01 |
| *walk* | 0.21 | 0.00 | 0.02 | 0.36 | 0.01 | 0.00 | 0.12 | 0.04 | 0.05 | 0.18 | 0.00 | 0.03 |
| *wd* | 0.06 | 0.00 | 0.00 | 0.00 | 0.00 | 0.00 | 0.22 | 0.00 | 0.00 | 0.00 | 0.72 | 0.00 |

**Table S7.** Transition probability from one behaviour (row name) to the other (column name) in the presence of a juvenile praying mantid. Transition probabilites are obtained by dividing each transition frequeny (see table S1) between a pair of behaviours by the total number of times a given behaviour was performed (row sums in table S1).

| Behavior | *ab* | *capture* | *fly* | *groom* | *jump* | *retreat* | *run* | *stop* | *occl* | *turn* | *walk* | *wd* |
| --- | --- | --- | --- | --- | --- | --- | --- | --- | --- | --- | --- | --- |
| *ab* | 0.00 | 0.01 | 0.05 | 0.01 | 0.01 | 0.05 | 0.06 | 0.03 | 0.02 | 0.03 | 0.73 | 0.00 |
| *capture* | 0.00 | 0.00 | 0.00 | 0.00 | 0.00 | 0.00 | 0.00 | 0.00 | 0.00 | 0.00 | 0.00 | 0.00 |
| *fly* | 0.00 | 0.00 | 0.00 | 0.06 | 0.00 | 0.01 | 0.04 | 0.48 | 0.02 | 0.00 | 0.40 | 0.00 |
| *groom* | 0.00 | 0.00 | 0.05 | 0.00 | 0.01 | 0.00 | 0.05 | 0.01 | 0.01 | 0.04 | 0.84 | 0.00 |
| *jump* | 0.00 | 0.00 | 0.00 | 0.03 | 0.00 | 0.00 | 0.00 | 0.41 | 0.00 | 0.00 | 0.56 | 0.03 |
| *retreat* | 0.39 | 0.00 | 0.09 | 0.00 | 0.00 | 0.00 | 0.00 | 0.04 | 0.04 | 0.30 | 0.13 | 0.00 |
| *run* | 0.08 | 0.00 | 0.02 | 0.14 | 0.00 | 0.00 | 0.00 | 0.13 | 0.01 | 0.09 | 0.52 | 0.00 |
| *stop* | 0.02 | 0.00 | 0.19 | 0.04 | 0.02 | 0.02 | 0.09 | 0.00 | 0.00 | 0.02 | 0.60 | 0.00 |
| *occl* | 0.01 | 0.00 | 0.03 | 0.08 | 0.01 | 0.00 | 0.05 | 0.01 | 0.00 | 0.00 | 0.80 | 0.00 |
| *turn* | 0.02 | 0.01 | 0.01 | 0.03 | 0.01 | 0.02 | 0.08 | 0.00 | 0.01 | 0.00 | 0.83 | 0.00 |
| *walk* | 0.15 | 0.00 | 0.05 | 0.34 | 0.02 | 0.01 | 0.05 | 0.14 | 0.08 | 0.16 | 0.00 | 0.05 |
| *wd* | 0.00 | 0.00 | 0.02 | 0.00 | 0.00 | 0.00 | 0.04 | 0.00 | 0.00 | 0.02 | 0.92 | 0.00 |

**Table S8.** Transition probability from one behaviour (row name) to the other (column name) before a juvenile praying mantid was introduced into the arena. Transition probabilites are obtained by dividing each transition frequeny (see table S1) between a pair of behaviours by the total number of times a given behaviour was performed (row sums in table S1).

| No mantid |  |  |  |  |  |  |  |  |  |  |  |  |
| --- | --- | --- | --- | --- | --- | --- | --- | --- | --- | --- | --- | --- |
| Behavior | *ab* | *capture* | *fly* | *groom* | *jump* | *retreat* | *run* | *stop* | *occl* | *turn* | *walk* | *wd* |
| *ab* | 0.00 | 0.00 | 0.00 | 0.00 | 0.00 | 0.03 | 0.03 | 0.00 | 0.00 | 0.03 | 0.91 | 0.00 |
| *capture* | 0.00 | 0.00 | 0.00 | 0.00 | 0.00 | 0.00 | 0.00 | 0.00 | 0.00 | 0.00 | 0.00 | 0.00 |
| *fly* | 0.00 | 0.00 | 0.00 | 0.03 | 0.00 | 0.00 | 0.00 | 0.59 | 0.03 | 0.00 | 0.34 | 0.00 |
| *groom* | 0.00 | 0.00 | 0.04 | 0.00 | 0.00 | 0.00 | 0.02 | 0.00 | 0.01 | 0.05 | 0.88 | 0.00 |
| *jump* | 0.00 | 0.00 | 0.00 | 0.21 | 0.00 | 0.00 | 0.07 | 0.29 | 0.00 | 0.00 | 0.43 | 0.00 |
| *retreat* | 0.33 | 0.00 | 0.00 | 0.00 | 0.00 | 0.00 | 0.00 | 0.33 | 0.33 | 0.00 | 0.00 | 0.00 |
| *run* | 0.00 | 0.00 | 0.00 | 0.35 | 0.04 | 0.00 | 0.00 | 0.04 | 0.00 | 0.00 | 0.58 | 0.00 |
| *stop* | 0.00 | 0.00 | 0.19 | 0.01 | 0.01 | 0.00 | 0.01 | 0.00 | 0.00 | 0.01 | 0.75 | 0.00 |
| *occl* | 0.00 | 0.00 | 0.02 | 0.03 | 0.00 | 0.00 | 0.00 | 0.00 | 0.00 | 0.02 | 0.93 | 0.00 |
| *turn* | 0.02 | 0.00 | 0.00 | 0.02 | 0.00 | 0.00 | 0.00 | 0.00 | 0.05 | 0.00 | 0.91 | 0.00 |
| *walk* | 0.07 | 0.00 | 0.02 | 0.50 | 0.02 | 0.00 | 0.04 | 0.10 | 0.10 | 0.15 | 0.00 | 0.04 |
| *wd* | 0.00 | 0.00 | 0.00 | 0.10 | 0.00 | 0.00 | 0.00 | 0.05 | 0.00 | 0.00 | 0.86 | 0.00 |

**Table S9.** Proportion of time spent in a given behavioural state by each individual fruit fly before and after introducing a treatment (i.e., a disturbance, spider or mantid) to the assay chamber.

| Treatment | Individual | Grooming |  | Walking |  | Running |  | Stopping |  | Retreat |  |
| --- | --- | --- | --- | --- | --- | --- | --- | --- | --- | --- | --- |
|  |  | Absent | Present | Absent | Present | Absent | Present | Absent | Present | Absent | Present |
| Disturbance | 1 | 0.820 | 0.728 | 0.180 | 0.265 | 0.000 | 0.005 | 0.000 | 0.000 | 0.000 | 0.000 |
|  | 2 | 0.590 | 0.265 | 0.338 | 0.658 | 0.000 | 0.009 | 0.044 | 0.015 | 0.000 | 0.000 |
|  | 3 | 0.264 | 0.219 | 0.658 | 0.685 | 0.048 | 0.036 | 0.013 | 0.052 | 0.000 | 0.000 |
| Spider | 1 | 0.528 | 0.490 | 0.359 | 0.394 | 0.022 | 0.011 | 0.000 | 0.003 | 0.000 | 0.000 |
|  | 2 | 0.116 | 0.000 | 0.844 | 0.861 | 0.000 | 0.086 | 0.000 | 0.053 | 0.000 | 0.000 |
|  | 3 | 0.706 | 0.181 | 0.262 | 0.368 | 0.010 | 0.076 | 0.000 | 0.347 | 0.005 | 0.002 |
| Mantid | 1 | 0.419 | 0.764 | 0.474 | 0.217 | 0.008 | 0.005 | 0.061 | 0.014 | 0.000 | 0.000 |
|  | 2 | 0.673 | 0.075 | 0.327 | 0.785 | 0.000 | 0.014 | 0.000 | 0.093 | 0.000 | 0.006 |
|  | 3 | 0.684 | 0.963 | 0.309 | 0.037 | 0.000 | 0.000 | 0.000 | 0.000 | 0.000 | 0.000 |

**Table S10.**
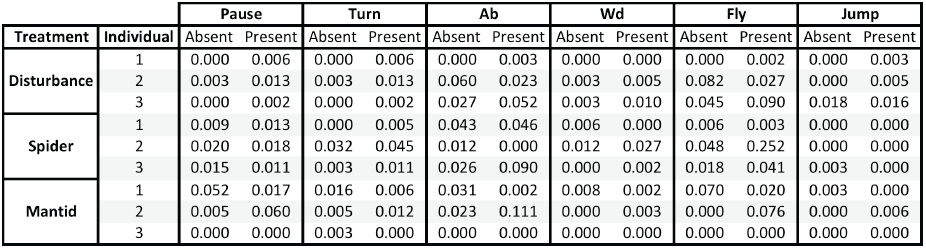
Number of occurrences per minute of each behavioural event before and after the introduction of a treatment (i.e., a disturbance, spider or mantid) to the assay chamber.

**Table S11.** Coefficients from a linear model (lm) for control individuals meansured before and after addition of a spider. While estimate of posterior means are similar to those of the main spider aset, Due to low sample sizes, CIs are large.

| *Grooming* |  |  |  |
| --- | --- | --- | --- |
| *Coefficient* | *Posterior mean* | *Lower CI* | *Upper CI* |
| Intercept | 181.88 | -23.34 | 387.10 |
| Predator state | -75.38 | -377.82 | 227.05 |
| Duration | 0.45 | -0.34 | 1.24 |
| *Locomotion* |  |  |  |
| Intercept | 156.76 | 37.32 | 276.21 |
| Predator state | 17.27 | -158.75 | 193.29 |
| Duration | 0.24 | -0.22 | 0.70 |

**Table S12.** Coefficients from a linear model (lm) for control individuals meansured before and er the addition of a mantid. Estimate of posterior means are similar to those of the main mantid aset, but due to low sample sizes, CIs large.

| *Grooming* |  |  |  |
| --- | --- | --- | --- |
| *Coefficient* | *Posterior mean* | *Lower CI* | *Upper CI* |
| Intercept | 134.79 | -1399.38 | 1668.96 |
| Predator state | 341.56 | -2661.10 | 3344.23 |
| Duration | -0.80 | -12.76 | 11.17 |
| *Locomotion* |  |  |  |
| Intercept | 345.69 | -919.99 | 1611.36 |
| Predator state | -323.02 | -2800.19 | 2154.15 |
| Duration | 1.64 | -8.24 | 11.51 |

**Table S13.** Coefficients from a linear model (lm) for control individuals meansured before and after isturbance. Despite low sample sizes, it is clear that disturbance had minimal effect on fruit fly haviours.

| *Grooming* |  |  |  |
| --- | --- | --- | --- |
| *Coefficient* | *Posterior mean* | *Lower CI* | *Upper CI* |
| Intercept | 200.00 | 60.11 | 339.81 |
| Disturbance | 0.00 | -197.78 | 197.78 |
| Duration | 2.38 | -0.53 | 5.29 |
| *Locomotion* |  |  |  |
| Intercept | 132.30 | -0.28 | 264.84 |
| Disturbance | 0.00 | -187.47 | 187.47 |
| Duration | -1.05 | -3.81 | 1.70 |

**Links to videos describing novel behaviours**

Abdominal Lifting http://dx.doi.org/10.6084/m9.figshare.1185638

Stopping Behaviour http://dx.doi.org/10.6084/m9.figshare.1185639

Retreat http://dx.doi.org/10.6084/m9.figshare.1185640

